# Novel Imaging Guidance for Cholesteatoma Surgery using Tissue Autofluorescence

**DOI:** 10.1101/2023.06.12.544537

**Authors:** Stella Yang, Joyce Farrell, Shenglin Ye, Iram Ahmad, Tulio A Valdez

## Abstract

**Significance:** Cholesteatoma is an expansile destructive lesion of the middle ear and mastoid, which can result in significant complications by eroding adjacent bony structures. Currently, there is an inability to accurately distinguish cholesteatoma tissue margins from middle ear mucosa tissue, causing a high recidivism rate. Accurately differentiating cholesteatoma and mucosa will enable a more complete removal of the tissue.

**Aim:** Develop an imaging system to enhance the visibility of cholesteatoma tissue and margins during surgery.

**Approach:** Cholesteatoma and mucosa tissue samples were excised from the inner ear of patients and illuminated with 405, 450, and 520 nm narrowband lights. Measurements were made with a spectroradiometer equipped with a series of different longpass filters. Images were obtained using a red-green-blue (RGB) digital camera equipped with a long pass filter to block reflected light.

**Results:** Cholesteatoma tissue fluoresced under 405 and 450 nm illumination. Middle ear mucosa tissue did not fluoresce under the same illumination and measurement conditions. All measurements were negligible under 520 nm illumination conditions. All spectroradiometric measurements of cholesteatoma tissue fluorescence can be predicted by a linear combination of emissions from keratin and flavin adenine dinucleotide (FAD). We built a prototype of a fluorescence imaging system using a 495 nm longpass filter in combination with an RGB camera. The system was used to capture calibrated digital camera images of cholesteatoma and mucosa tissue samples. The results confirm that cholesteatoma emits light when it is illuminated with 405 and 450 nm, whereas mucosa tissue does not.

**Conclusions:** We prototyped an imaging system that is capable of measuring cholesteatoma tissue autofluorescence.

## 1. INTRODUCTION

Cholesteatoma is a proliferative mass of keratinized epithelium overlying a fibrous stroma that occurs in the middle ear and mastoid cavity.^1^ The term “cholesteatoma” was first used in 1838 by Johannes Müller (1801-1858), a German scientist and author most notable for his work in psychology and pathology,^2^ based on his belief that the mass was composed mostly of cholesterine.^3^ Despite the term cholesteatoma being a misnomer (as this benign mass contains neither cholesterine nor fat), it is still the most commonly used term to refer to these masses. It is widely recognized that cholesteatomas are divided into two categories: congenital and acquired. Cholesteatomas are considered congenital when it presents in the absence of prior surgery or perforation and usually occur in the pediatric population.^4, 5^ These lesions are often asymptomatic and are detected either during routine pediatric visits or when the disease is grossly advanced. When cholesteatoma cases become symptomatic, common manifestations include: otorrhea,^6^ otalgia,^7^ and hearing loss.^8^ Acquired cholesteatoma occurs at an annual rate of about 9-14 cases per 100,000 adults, and 3-17 cases per 100,000 children. Furthermore, there is a greater amount of acquired cholesteatoma, roughly 70-96% of all cases

If left untreated, cholesteatomas can lead to severe complications such as brain abscesses, meningitis, facial nerve paralysis, and hearing loss.^9^ There is no medical treatment for cholesteatoma and it requires complete surgical removal of the mass via tympanomastoidectomy to prevent recidivism.^10, 11^ Unfortunately, cholesteatomas recur with rates ranging from 5 to 50% thereby necessitating multiple surgical procedures.^12, 13^ One of the principal determinants of this high rate of recidivism is residual cholesteatoma left behind after the surgery – a direct consequence of the inability to accurately visualize the mass margins due to its difficulty of being differentiated from healthy, mucosa tissue.^14^ Thus, there is an unmet need for an imaging system to augment the real-time diagnostic information available to the surgeon in order to enable more complete removal of the mass.^15^ The imaging system must also provide the benefits of the current gold standard of care such as being able to inspect of all quadrants of the tympanic membrane and middle ear, and being specific enough to distinguish between other similar masses behind the tympanic membrane from those within (e.g., myringosclerosis).^16^

The capability of performing an accurate assessment in congenital cholesteatomas is further compromised by difficulties in examining the middle ear in pediatric patients.^17^ Potential pitfalls include the presence of narrow external auditory canals and an uncooperative child. Despite these critical challenges, the otoscope, which is the most common instrument for middle ear examination, has undergone few changes since the mid-19th century.^18^ Otoscopic examination still relies on light reflecting from the tympanic membrane and interpretation of the findings by the physician. Taken together, these visualization challenges make the diagnosis of a congenital cholesteatoma difficult for both pediatricians and otolaryngologists. Similar challenges are encountered in the operating room where white light and magnification using a microscope or endoscopes have not resulted in decreased cholesteatoma recurrence. In this context, the ability to obtain biochemical information of the tympanic membrane and middle ear masses would provide a much-needed dimension to the diagnosis of cholesteatoma.

Our current research seeks to combine the molecular basis of spectroscopy with wide-field imaging capabilities that could potentially serve to bridge the chemical and morphological domains. The spectroscopic measurement of tissue chemistry for middle ear examination is, however, largely unexplored and no analytical method exists to reproduce or exceed otoscopic diagnostic capabilities. Based on our experience in elucidating latent information about this diverse range of pathophysiological conditions, we hypothesize that the utility of such spectroscopic imaging can be extended to identify these critical structures in the middle ear due to their unique chemical composition in relation to the neighboring tissue.

Cholesteatoma is known to be largely comprised of keratin debris that is surrounded by a thin layer or matrix of stratified squamous epithelium and and an outer layer or perimatrix of fibrous tissue.^19–21^ Therefore, it would be beneficial to design an imaging system that can take advantage keratin’s fluorescent properties to excite and measure its fluorescence. However, the peak excitation wavelength for keratin (278—279 nm) would also damage healthy tissue.^22^ Consequently, it behooves us to search for excitation lights that do not damage healthy tissue and, at the same time enhance the visibility of cholesteatoma tissue relative to healthy, mucosa tissue.^23^

Preliminary evidence suggests that narrowband illumination with peak spectral energy at 405 and 450 nm can excite keratin in epithelial tissue extracted from rabbits and humans.^24^ To investigate this possibility, our research efforts have focused on the acquisition of fluorescence images based on a modified surgical endoscope.^25^ This novel otoscope offers multi-wavelength, video-rate fluorescence imaging and provides a first step in evaluating this new approach for diagnosis of middle ear pathology. The underlying basis of these measurements, however, remains poorly understood. In particular, spectroscopic measurement of the tissue biochemistry and its sensitivity to localized changes requires systematic investigation. In this article, we aim to describe the *in vivo* label-free fluorescence data acquired using this endoscope in the context of its chemical composition. Here we employ intrinsic autofluorescence spectroscopy by exciting the constituents as well as the sample with 405 nm and 450 nm light.^26^ The 520 nm light was also used as a control. Based on the spectral signatures of the chemical-morphological model constituents, the tissue spectral information acquired is then translated to provide quantitative measures of the chemical makeup of the lesion interrogated. By correlating these results with fluorescence microscopy and histopathology observations, it may be possible to determine differential molecular markers in tissue and in disease. We expect that this determination will not only allow better visualization of lesion margins but also provide specific molecular insight into disease progression. Such automated spectroscopic recognition potentially provides a route to accurate, reproducible, and economical diagnosis without depending solely on human visual interpretation.

## 2. MATERIALS AND METHODS

### 2.1 Spectroradiometer Device

Cholesteatoma and mucosa (normal, healthy middle ear tissue) samples were excised from patients and illuminated with different narrowband lights. The light source was a customized Sony laser diode video scope illumination unit (model #MBURD-RGBW-5G, S/N002) capable of creating narrow band and broadband illumination, including IR and white light. Figure 1a plots the spectral energy in the 3 narrowband lights, with peak energy at 405, 450 and 520 nm, that were used in this study. The light source was placed 30 mm from the tissue surface and the power was set to 40 mW.

**Figure 1.**
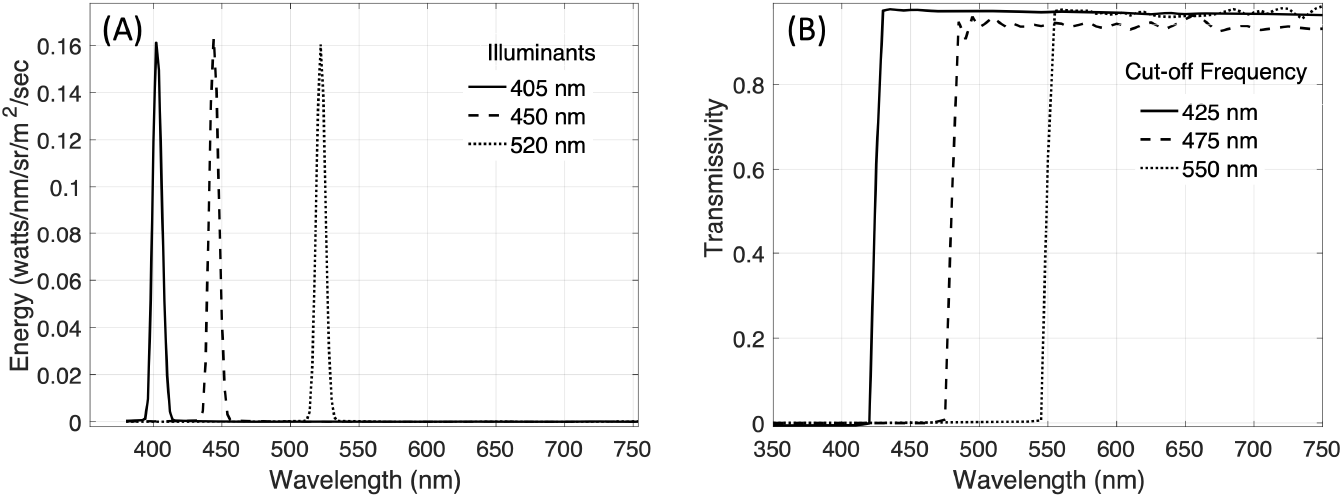
Spectral energy of narrow-band illumination and spectral transmissivities of the long pass filters. (A) The spectral energy of lights with peak energy at 405, 450 and 520 nm are plotted in solid, dashed and dotted lines, respectively. (B) The spectral transmissivity of the 425, 475 and 550 nm long pass filters are plotted in solid, dashed and dotted lines, respectively.

All spectral measurements were made with a SpectraScan® PR-670 spectroradiometer with a 1-degree aperture. We used long pass filters: Edmund Scientific High Performance OD4 Longpass Filters, stock Numbers 84748 (425 nm 50 mm), 84749 (475 nm, 50 mm) and 94757 (550 nm, 50 mm), to attenuate the laser illumination. One of the three long pass filters was placed over the aperture of the PR-670 in order to block the excitation light from reaching the sensor. Calibration measurements confirmed that light reflected from the illumination light was blocked by all the relevant filters. The spectroradiometric measurements reported in the results section below were obtained 1) when the tissue was illuminated with a 405 nm light and the PR-670 was equipped with a 425 nm longpass filter, 2) when the tissue was illuminated with a 450 nm light and the PR-670 was equipped with a 475 nm longpass filter, and 3) when the tissue was illuminated with a 520 nm light and the PR-670 was equipped with a 550 nm longpass filter. Figure 1b plots the spectral transmissivity of the 425, 475 and 520 nm longpass filters. spectroradiometric measurements of keratin and flavin adenine dinucleotide (FAD) powder were also obtained under the same illumination and filter conditions.

### 2.2 Imaging Device

Images of the illuminated tissue samples were obtained using an Allied Vision 1800 U-319C red-green-blue (RGB) color camera with a Thor Labs FGL495 - Ø25 mm GG495 Colored Glass Filter, 495 nm longpass filter. The camera was placed 10 cm from the sample at an angle that was normal to the surface of the tissue and the illumination conditions were the same as the illumination conditions for the spectroradiometric measurements. The camera images were obtained using an exposure duration of 500 ms in order to capture camera images that had the highest signal-to-noise ratio (SNR) and did not include saturated pixel values.

### 2.3 Microscope Slides

Cholesteatoma and mucosa sample tissues were embedded in Tissue-Tek® O.C.T. Compound and cryosectioned at -20°C into a thickness of 10 μm, then placed onto glass slides for autofluorescence imaging. The microscope images were taken by a ZEISS LSM700 using its 405 nm and 488 laser diode, both at 5mW of power. The microscope’s 10x objective (model#: EC Plan-Neofluar 420340-9901), and PMT fluorescence detector were used. Corresponding H&E stained images were also viewed under a bright-field microscope with a 10x objective. All imaging data is displayed using Fiji - ImageJ.

## 3. RESULTS

### 3.1 Spectroradiometric Measurements

Figure 4a plots the PR-670 spectroradiometric measurements that were made when several different cholesteatoma tissue samples were illuminated with a 405 nm light in combination and a 425 nm longpass filter. Figure 4b shows the spectroradiometric measurements for the same tissues when they were illuminated with the 450 nm light and measured by the PR-670 through a 475 nm longpass filter. And Figure 4c shows the measurements made when the tissues were illuminated with the 520 light and measured with the PR-670 through a 550 nm filter.

**Figure 2.**
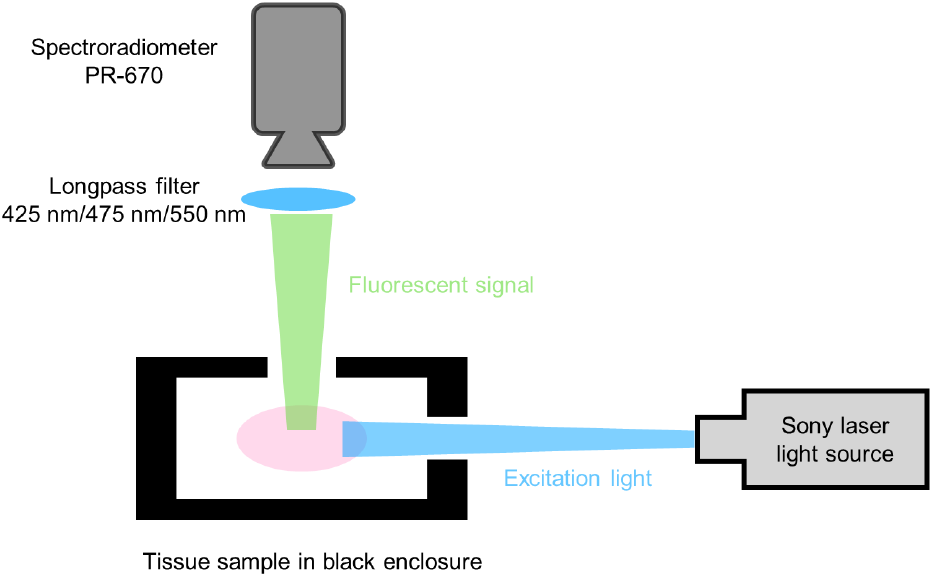
The customized Sony (model #: MBURD-RGBW-5G, S/N002) laser diode video scope illumination unit with 3 different wavelengths (405nm, 450nm and 520nm @ 40W output) was applied as the illumination source to excite the tissue sample which was secured in the black enclosure to avoid the ambient light reflection. Longpass filters of 425 nm, 475 nm, and 550 nm (Edmund Scientific High Performance OD4 Longpass filter 84748, 84749 and 94757, respectively) were used to block the excitation wavelength and any other reflected light. The spectroradiometer (Photon research 670) was applied to measure the fluorescent spectra.

**Figure 3.**
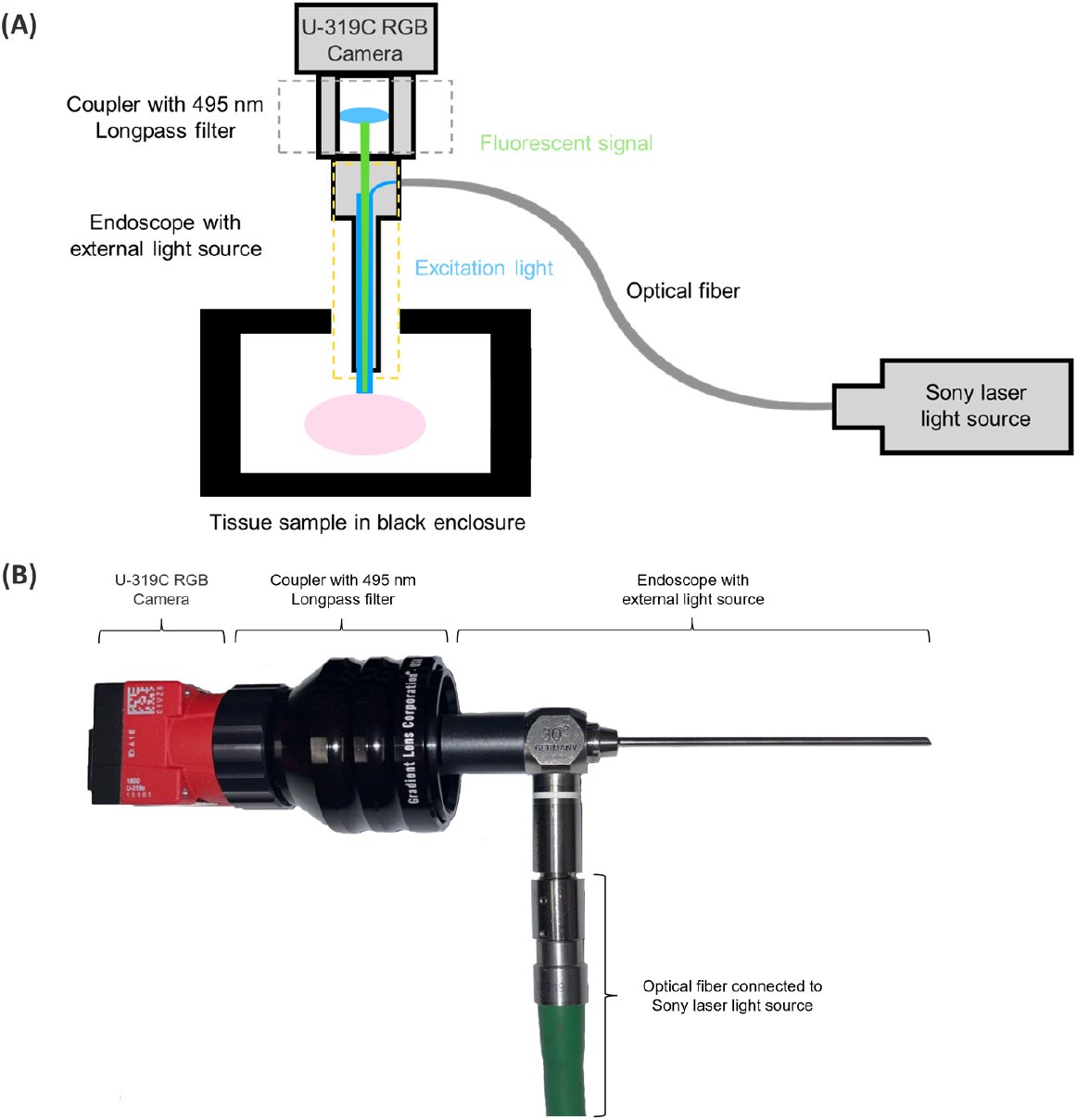
The tissue sample was secured in the black enclosure to avoid any ambient light reflection. An endoscope equipped with an external light source (Sony laser diode video scope illumination unit) was applied to excite and collect the fluorescence signal from the tissue sample. The U-319C RGB camera was used as the image sensor to capture the fluorescence image. For middle ear access, we used a Karl-Storz Tele-Otoscope (model 1218 AA, color code: green) with HOPKINS Straight Forward Telescope 0^*°*^, diameter 3 mm, length 6 mm, with a magnification of 4X for a field of view of about 1 cm by 1 cm. An optical coupler was used to connect the endoscope and camera to avoid light leakage. The 495 nm longpass filter was embedded into the optical coupler and placed in front of the camera to block the excitation wavelength and any other reflected light. (A) Diagram of the setup (B) Actual imaging device.

**Figure 4.**
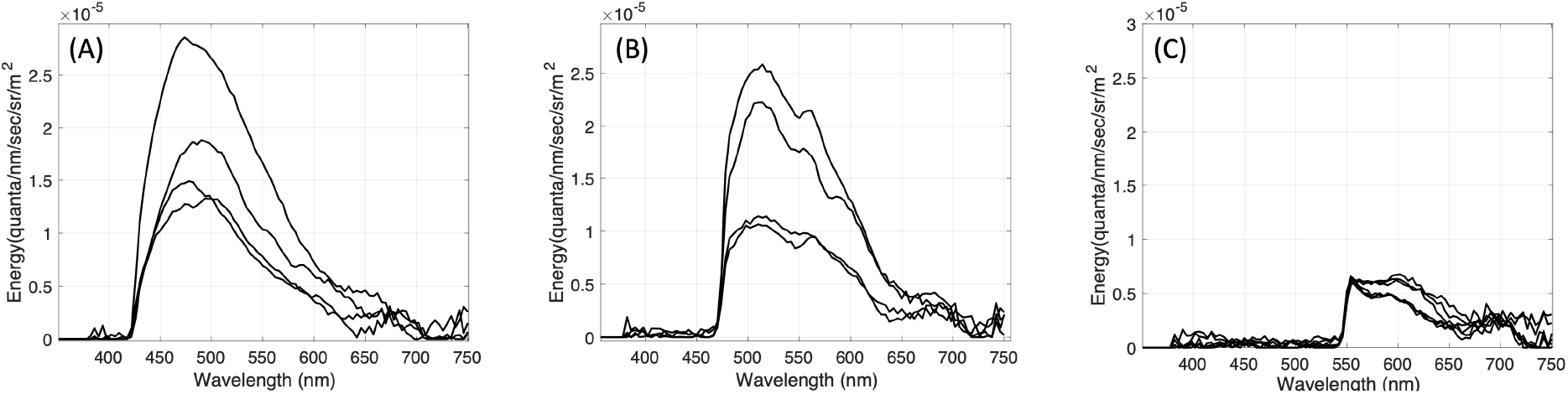
Cholesteatoma tissue fluorescence measurements. Different tissue samples were illuminated with narrowband light (see Figure 1a). Fluorescence measurements (reported in units of radiance) were made using a spectroradiometer equipped with longpass filters (see Figure 1b) to block reflected light. (A) spectroradiometric measurements obtained when cholesteatoma tissue samples were illuminated with the 405 nm light and reflected light was blocked with the 425 nm longpass filter. (B) Measurements obtained with the 450 nm light and the 475 nm longpass filter. (C) Measurements obtained with the 520 nm light and the 550 nm longpass filter.

To confirm that the spectroradiometric measurements shown in Figure 4 are the result of tissue autofluorescence and not tissue reflectance, we estimated the amount of light that we would expect to be reflected from cholesteatoma tissue. This estimate was obtained in the following way. First, we multiplied the spectral energy of each light with the filter transmittance of the longpass filter. This estimates the maximum spectral energy in each light that could be reflected from a diffusely reflecting surface. To estimate the amount of light that could be reflected from cholesteatoma tissue, we multiplied the maximum reflected light with the measured cholesteatoma tissue reflectance. (Tissue reflectance measurements are based on spectrophotometric measurements of cholesteatoma tissue illuminated by a halogen lamp divided by the spectrophotometric measurements of a white calibration surface also illuminated by the halogen and placed in the same position as the cholesteatoma tissue.) Figure 5 shows that the measurements of cholesteatoma tissue autofluorescence for the 405 nm light is greater than the light that could be reflected from cholesteatoma tissue.

**Figure 5.**
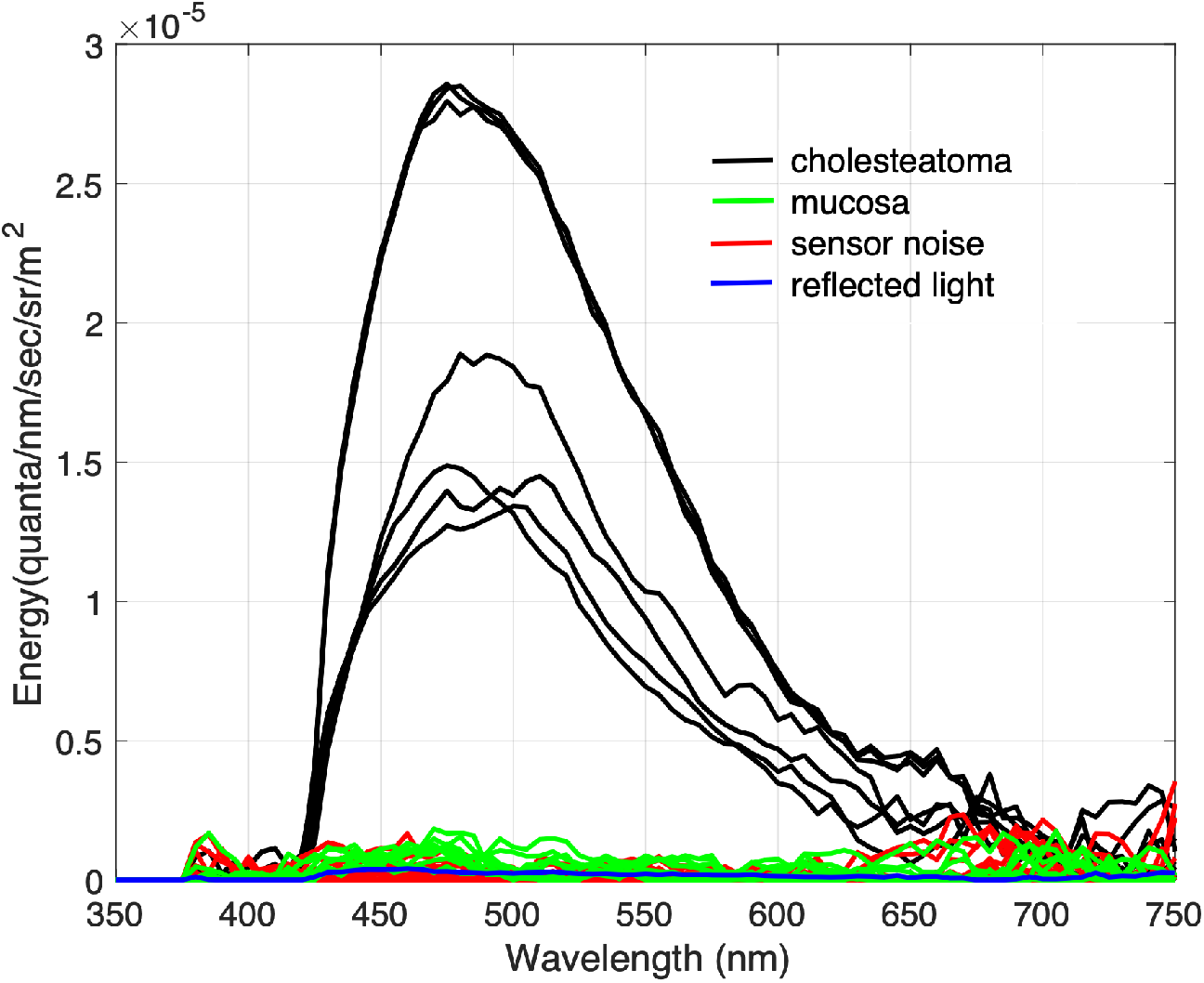
Comparison of tissue fluorescence, sensor noise and estimated tissue reflectance. The spectroradiometric measurements were obtained when cholesteatoma tissue samples (plotted in black), mucosa tissue samples (plotted in green) and black fabric (plotted in red) were illuminated (separately) with the 405 nm light and reflected light was blocked with the 425 nm longpass filter. The blue lines plot the maximum possible reflected light estimated by multiplying the product of the spectral power of the 405 nm light and the spectral transmittance of the 425 nm longpass filter with measured cholesteatoma tissue reflectance..

We also compared the spectroradiometric measurements of tissue autofluorescence with the spectroradiometric measurements of a black surface under the same illumination and filter conditions. In the absence of reflected light, spectroradiometric measurements of the black surface quantify the noise in the spectroradiometer itself. Figure 5 shows that our measurements of cholesteatoma tissue autofluorescence are greater that measurements of light that may be reflected from cholesteatoma tissue and greater than measurements of sensor noise.

Figure 5 also shows that the spectroradiometric measurements of mucosa tissue are similar to the measurements of reflected light and sensor noise. From these data, we conclude that the samples of mucosa tissue did not fluoresce when illuminated by any of the excitation lights, whereas the samples of cholesteatoma did fluoresce when illuminated with 405 and 450 nm lights.

### 3.2 Fluorophore Predictions

In order to better understand the fluorescence signal we are measuring, we opted to deconstruct the signal into its components. We tested compounds such as Nicotinamide adenine dinucleotide (NADH),^27, 28^ FAD,^29^ elastin,^30^ flavin,^31^ protoporphyrin IX,^32^ collagen,^33, 34^ and keratin.^35, 36^

Figure 6A plots spectroradiometric measurements obtained when keratin powder was 1) illuminated with the 405 light and filtered with the 425 nm longpass filter (solid line) and 2) illuminated with the 450 light and filtered with the 475 nm longpass filter (dashed line). The measurement data are consistent with excitation emission matrices for keratin powder that have been published in the literature.^37^ For example, keratin is known to have broad excitation and emission spectra with peaks at 275 nm and 375 nm, respectively.^38^ Other studies have reported that a 405 nm light is more effective at exciting keratin fluorescence than a 457 nm light.^35^ Our measurements of keratin fluorescence are consistent with these prior observations. Although the peak of the measured emission spectra shown in Figure 6a appear to shift, this effect is due to the fact that the longpass filters block both emitted and reflected light. The 425 nm longpass filter is necessary to block light that is reflected from the 405 nm light, but it also blocks light that is emitted by keratin powder below 425 nm as well. Similarly, the 475 nm longpass filter blocks light that is reflected from the 450 nm light, but also blocks light that is emitted by keratin powder below 475 nm. Hence, the apparent shift in the peak of the measured fluorescence spectra for keratin powder is due to the effect of the longpass filters and not due to any change in the excitation emission properties of keratin powder. The difference in the amplitude of the fluorescence emissions is due to the fact that 405 is a more effective excitation wavelength for keratin fluorescence.

**Figure 6.**
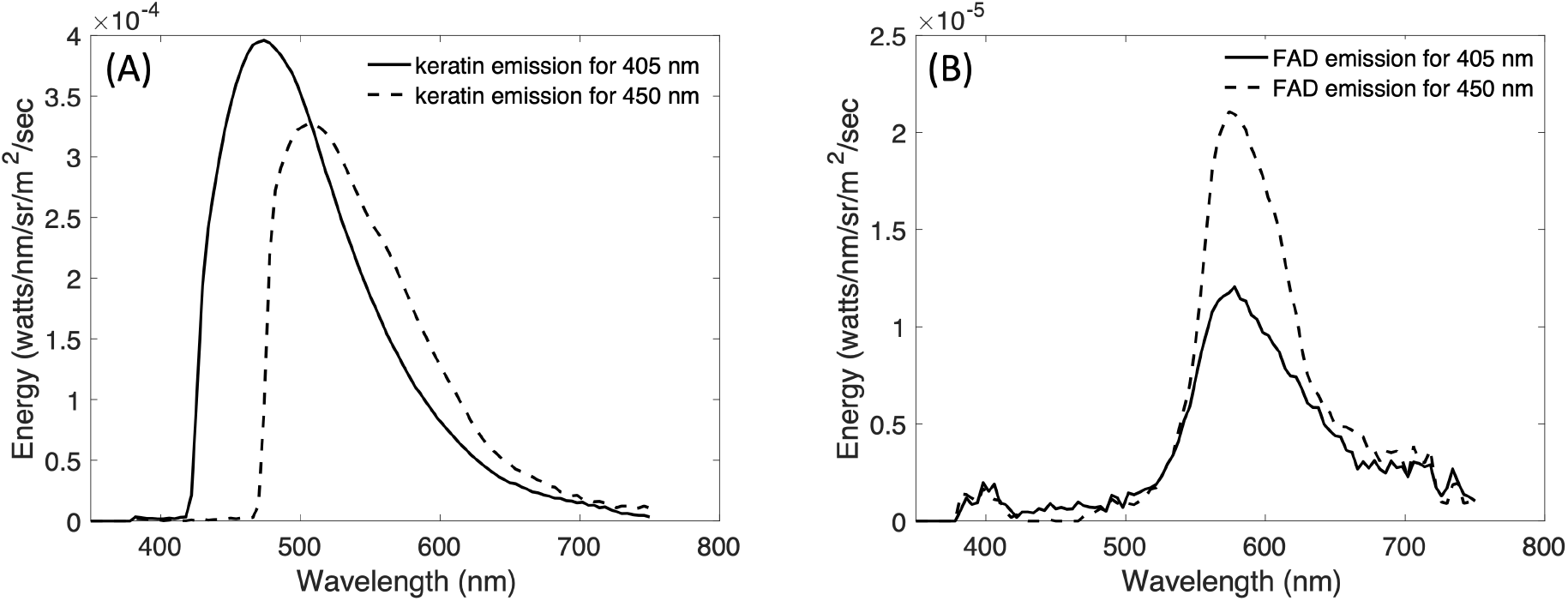
Keratin and FAD fluorescence measurements. (A) The spectroradiometric measurements of keratin powder illuminated with the 405 nm light and filtered with the 425 nm longpass filter is plotted as a solid line and measurements of the same keratin powder illuminated with the 450 light and filtered with the 475 nm longpass filter is plotted as a dashed line. (B) The spectroradiometric measurements of FAD powder illuminated with the 405 nm light and filtered with the 425 nm longpass filter is plotted as a solid line and measurements of the same FAD powder illuminated with the 450 light and filtered with the 475 nm longpass filter is plotted as a dashed line.

Figure 6b plots spectroradiometric measurements obtained when FAD powder was 1) illuminated with the 405 light and filtered with the 425 nm longpass filter (solid line), and 2) illuminated with the 450 light and filtered with the 475 nm longpass filter (dashed line). These measurements are also consistent with published and well-established measurements of the excitation and emission properties of FAD.^37^ Fluorescence emissions peak around 560 nm for both the 405 and 450 nm lights, hence the 425 and 475 nm long pass filters do not block any of the emitted light. The difference in the amplitude of the fluorescence emissions is due to the fact that 450 is a more effective excitation wavelength for FAD fluorescence.

We used the spectroradiometric measurements shown in Figure 6 of keratin and FAD as spectral basis functions to test the hypothesis that the cholesteatoma tissue fluorescence measurements for each light can be predicted by a linear combination of keratin and FAD emissions measured for the same lighting conditions.

We used non-negative least-squares regression to test the hypothesis that the cholesteatoma tissue fluorescence can be explained by a weighted combination of spectral basis functions (emissions) from keratin and FAD (see Figure 6). Figure 7 plots the predicted and measured spectroradiometric measurements of cholesteatoma tissue fluorescence measured for 4 different tissue samples, each illuminated by the 405, 450 and 520 nm excitation lights. The predictions are based on a weighted combination of the keratin and FAD emissions. In all cases, the predictions using a non-negative least-squares regression account for more that 94% of the variance in the measurements.

**Figure 7.**
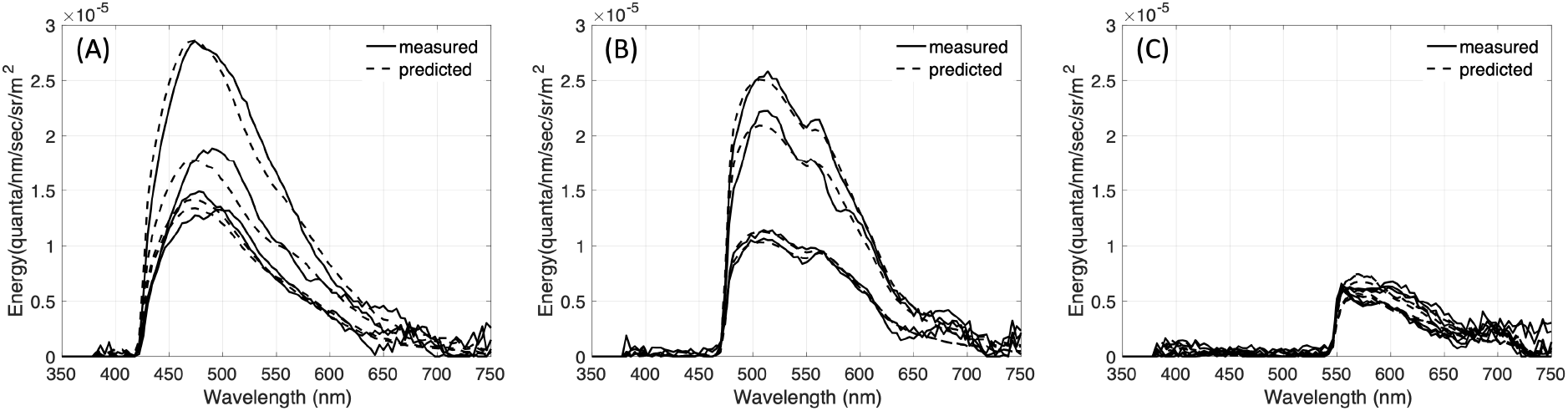
Cholesteatoma tissue fluorescence (solid lines) can be predicted by a linear combination of keratin and FAD fluorescence (dashed lines). (A) Measured and predicted cholesteatoma tissue fluorescence excited by the 405 nm light. (B) Measured and predicted cholesteatoma tissue fluorescence excited by the 450 nm light. (C) Measured and predicted cholesteatoma tissue fluorescence excited by the 520 nm light.

### 3.3 Imaging Device Measurements

Figure 8 shows U-319C camera images of cholesteatoma and mucosa tissue reflectance and fluorescence. Reflectance images were obtained by illuminating the tissue with a white light and capturing camera images with no filter in place. Fluorescence images were obtained by illuminating the tissue with 405 nm narrowband light or with 450 narrowband light and capturing camera images with a 495 nm long pass filter in place. The camera images illustrate that it is possible detect fluorescence in the cholesteatoma tissue samples but not in the mucosa tissue samples. This result is consistent with the spectroradiometric measurements of cholesteatoma and mucosa tissue that are shown in Figure 5. Since the mucosa tissue does not fluoresce under 405 or 450 nm light, the camera images of mucosa tissue should be black, as they are.

**Figure 8.**
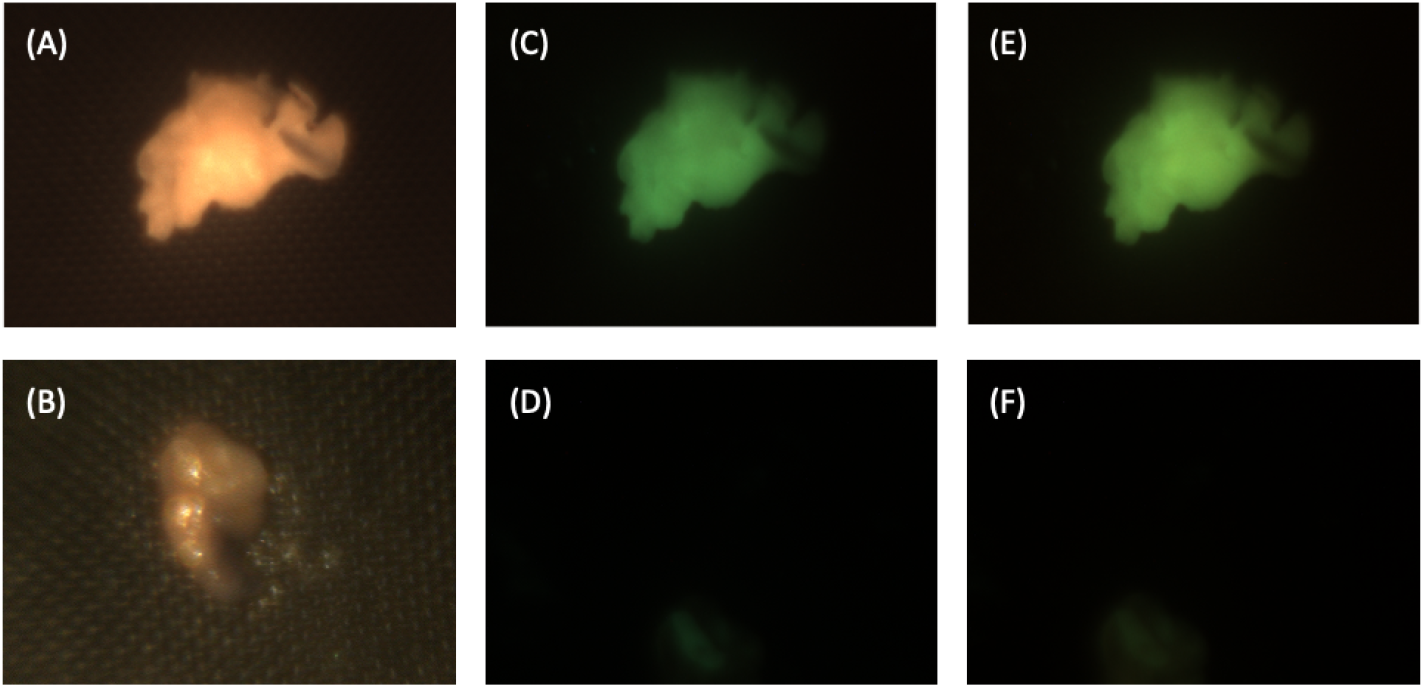
Reflectance and fluorescence images of cholesteatoma and mucosa tissue samples. To visualize tissue reflectance, the cholesteatoma (A) and mucosa (B) tissue samples were illuminated with white light and captured with no filter. Tissue fluorescence images were measured when the same cholesteatoma (C) and mucosa (D) tissue samples were illuminated with the 405 nm light and captured with a 495 nm longpass filter on the U-310c digital camera, using a 500 ms exposure duration. Tissue fluorescence images were also measured when the same cholesteatoma (E) and mucosa (F) tissue samples were illuminated with the 450 nm light and captured with a 495 nm longpass filter on the camera and a 500 ms exposure duration.

Since the fluorescence signal intensity is higher when cholesteatoma tissue is illuminated with 405 nm light than with 450 nm light, one might expect the camera images obtained when the tissue is illuminated with 405 nm light to be brighter than the images obtained when the tissue is illuminated with 450 nm light. This was not what we observed, however. Figure 9 shows camera images for two additional cholesteatoma samples. As seen in Figure 8, the camera images captured when cholesteatoma tissue is illuminated with 450 nm light appear to be brighter than the camera images captured when the tissue is illuminated with 405 nm light. We calculated the mean R, G and B pixel values for a bright region in the camera images obtained when cholesteatoma tissue samples were illuminated by 405 and 450 nm light with a fixed exposure duration (500 ms). The mean values are dependent on the region selected, but when we consistently chose the same region in the camera images of each tissue sample, the pixel values obtained under 450 nm light illumination were consistently higher compared to the pixel values obtained under 405 nm.

**Figure 9.**
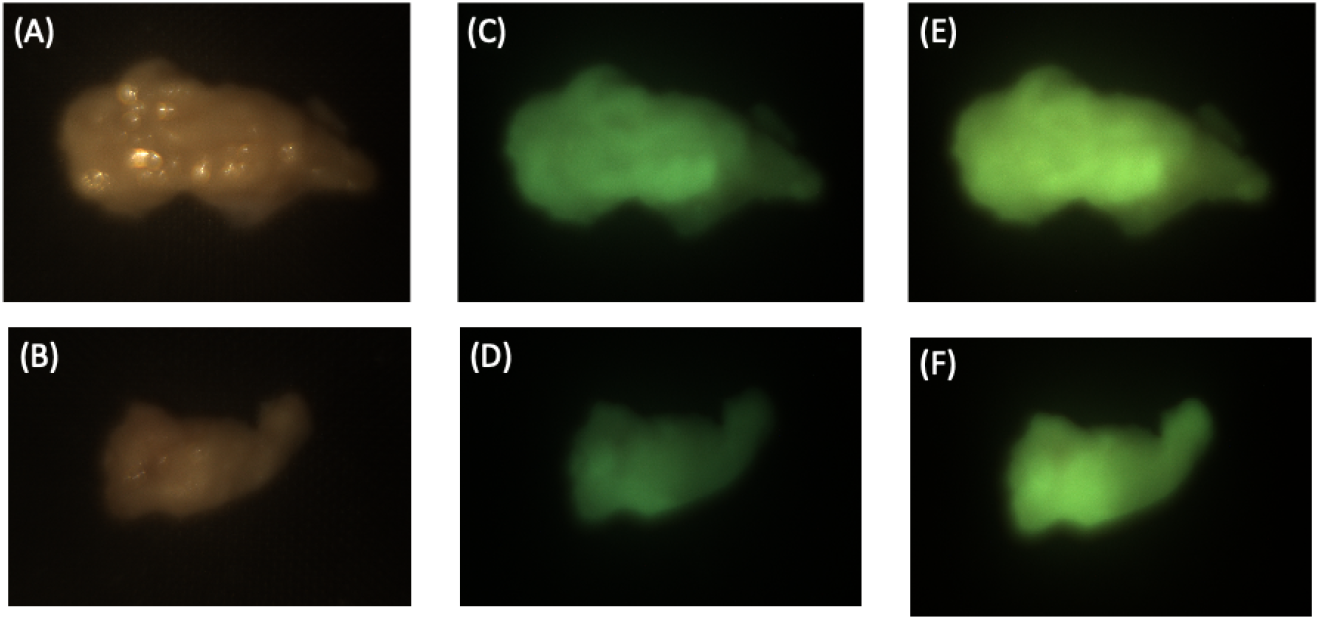
U-319C camera images of 2 additional samples of (A and B) cholesteatoma tissue illuminated with white light. Cameras images of cholesteatoma tissue (C and D) illuminated with 405 nm light and captured with a 495 nm longpass filter and (E and F) illuminated with 450 nm light and captured with a 495 nm longpass filter.

This seemingly paradoxical result can be explained by analyzing the spectral quantum efficiency (QE) of the camera. Figure 10 compares the spectrophotometric measurements of cholesteatoma tissue fluorescence with the camera quantum efficiency (QE) (calculated by multiplying the spectral transmittance of the 495 nm longpass filter with the published spectral quantum efficiency of the camera’s imaging sensor). The overlap between the camera QE (specifically, the green color channel of the imaging sensor) and the signal (specifically the cholesteatoma tissue fluorescence) is greater when tissue is illuminated with 450 nm light than when it is illuminated with 405 nm light. The qualitative comparison of the overlap of the signal and sensor shown in Figure 8 is supported by a quantitative analysis of the signal strength and camera QE.

**Figure 10.**
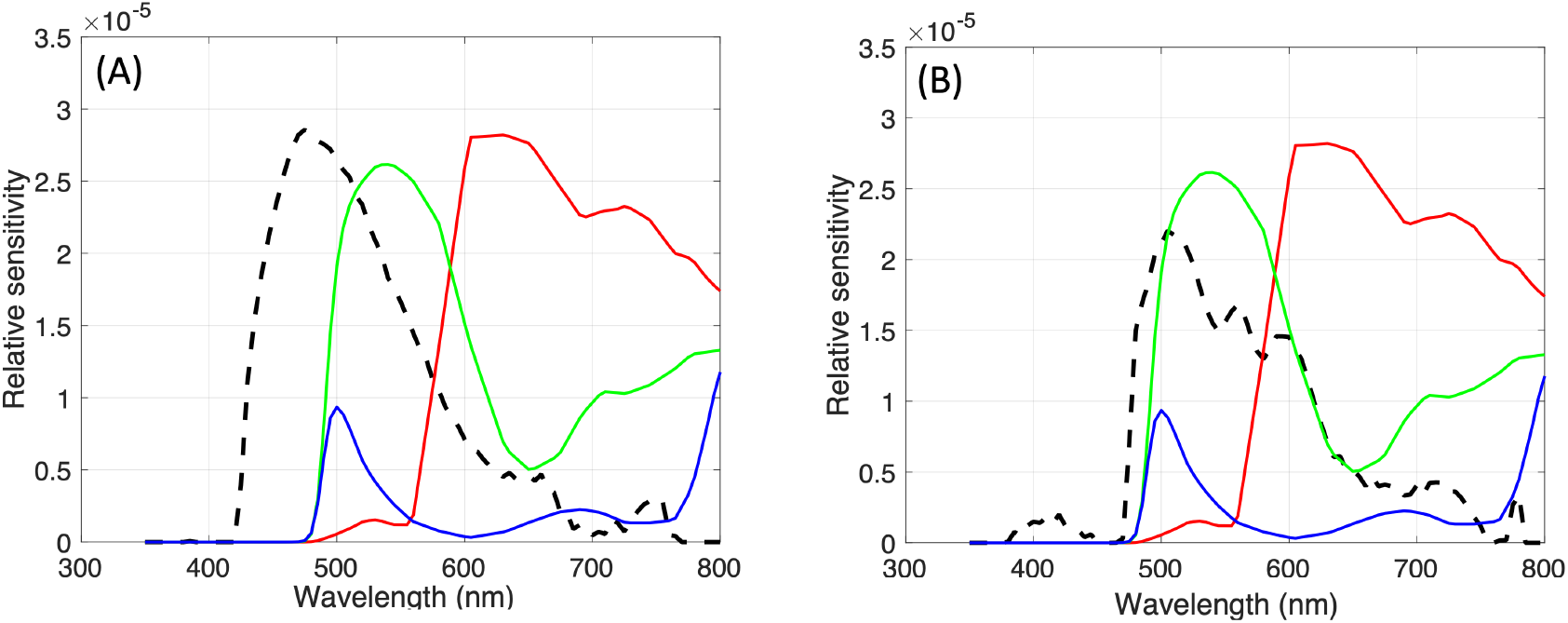
Comparing the signal (dashed lines) with the sensor (solid red, green and blue lines). A) The spectral radiance measured when a cholesteatoma tissue sample was illuminated with 405 nm light is plotted as a dashed line and the quantum efficiency (QE) of the 3 color channels in the digital camera are plotted as solid lines. To facilitate comparisons, the camera QE (a product of the sensor spectral sensitivities and the 495 nm longpass filter spectral transmittance) has been scaled to match the maximum spectral radiance measurements. B) The spectral radiance measured when the same cholesteatoma tissue sample was illuminated with 450 nm light (dashed line) is compared to the scaled camera QE (solid lines).

The camera pixel values for each illumination condition can be predicted by multiplying the camera QE with the spectrophotometric measurements. More specifically, the camera QE can be expressed as a 3xW matrix, S, where W refers to the number of wavelength samples. The spectrophotometric readings of cholesteatoma tissue fluorescence under excitation from either 405 or 450 nm light is represented as a vector, F, of length W. The camera RGB values, R, are obtained by multiplying the matrix S and vector F (*R* = *S · F*). We evaluated the measured and predicted camera pixel values for cholesteatoma tissue illuminated by both 405 and 450 nm lights. The green (G) pixel values, both measured and predicted, were found to be higher than the red (R) and blue (B) pixel values, and the predicted and measured G values were higher for the 450 nm light than for the 405 nm light. The alignment of predicted and measured camera pixel values supports our understanding of the effect of the camera components.

### 3.4 Microscope Slides

Figure 11 shows microscope slides of 10 μm in thickness of a cholesteatoma keratin pearl and mucosa tissue under different fluorescent conditions compared to a hematoxylin and eosin (HE) stained condition. Under 405 and 488 light illumination, the mucosa tissue sample shows the lack of autofluorescence which is consistent with the macro-level spectroradiometric and camera images. Under the same illumination, the keratin pearl cholesteatoma sample also shows consistent autofluorescence properties. The fluorescence on the micro-scale offers further insight into the detailed distribution of the emitted fluorescence light. The center of the pearl, which contains the most keratin, also emits the most fluorescence, relative to the surrounding cholesteatoma tissue. This could indicate that keratin could be responsible for more of the fluorescence signal on the macrolevel, or perhaps locations where there are more keratin correlate with locations where there are greater levels of FAD as well. Looking at the merged images (Figure 11c and 11g), while the mucosa seems to show homogeneous distribution of fluorescence under 405 and 450 nm illumination, the keratin pearl has certain areas where one excitation wavelength shows stronger fluorescence over the other, which highlights the benefits of using both wavelengths to illuminate different fluorophores. Overall, the microscope slides are consistent with the images acquired from the imaging device as well as the spectral measurements recorded by the spectroradiometer.

**Figure 11.**
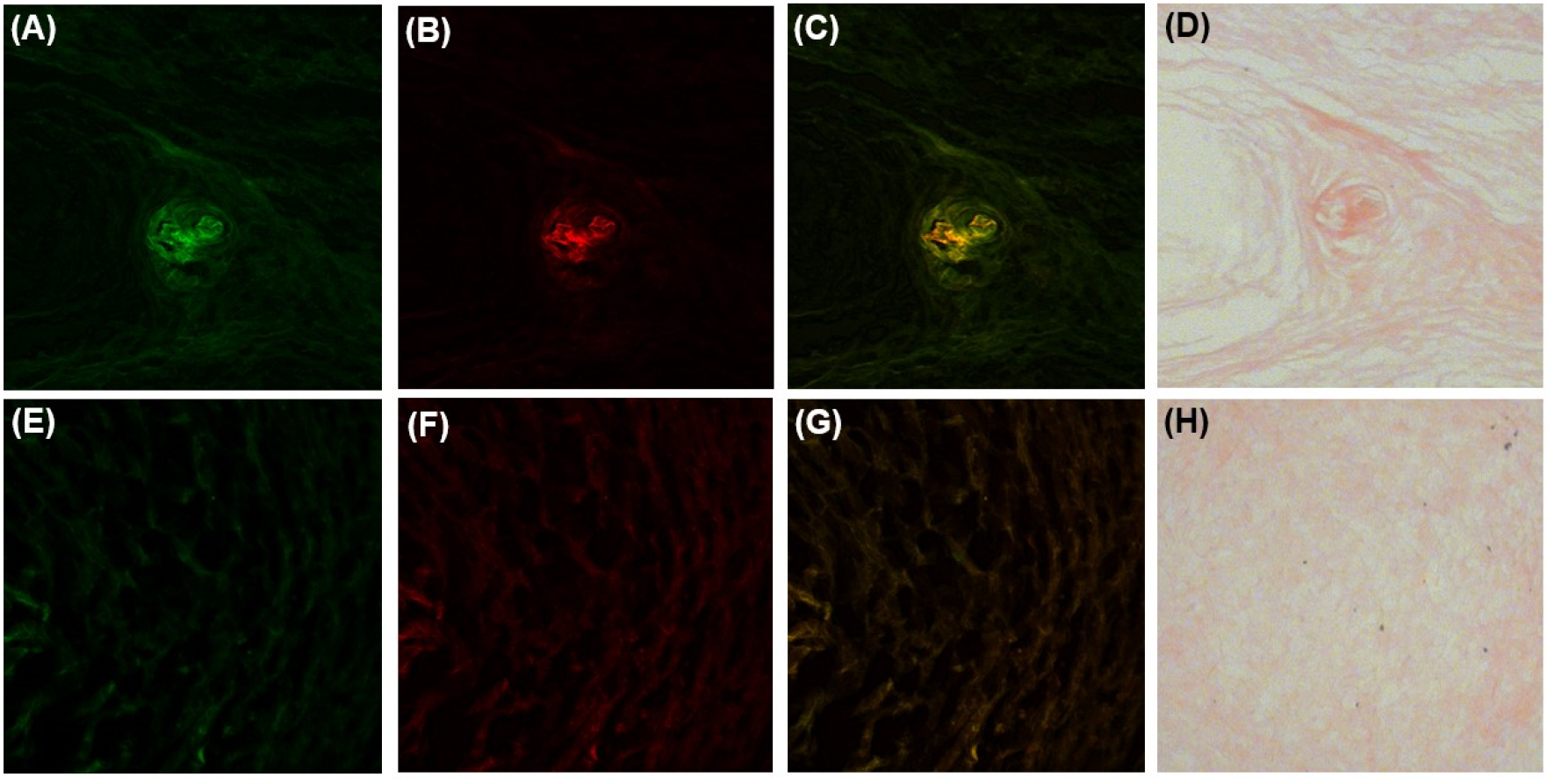
Autofluorescence slide images of cholesteatoma keratin pearl sample and mucosa sample (A and E) illuminated under 405 nm light, (B and F) illuminated under 488 light, (C and G) merged with 405 nm light represented in green and 450 nm light represented in red, and (D and H) stained using hematoxylin and eosin.

## 4. DISCUSSION

Recurrence following surgical excision continues to be a challenge in the management of cholesteatoma.^39^ The biggest reason for this issue is the difficulty of distinguishing cholesteatoma from surrounding mucosa and granulation tissue. In this study, we obtained cholesteatoma and mucosa autofluorescence from surgically removed samples from patients and successively illuminated with 405, 450, and 520 nm narrowband lights. Spectral measurements of the tissue were made using a spectroradiometer equipped with 425, 475, and 550 nm longpass filters that blocked reflected light from the 405, 450, and 520 nm illumination conditions, respectively. The spectroradiometric measurements showed significant cholesteatoma tissue autofluorescence under 405 nm and 450 nm illumination, as well as a lack of fluorescence from mucosa tissue. The fluorescence of tissue illuminated at 520 nm was extremely low and almost indistinguishable from sensor noise. We show that it is possible to predict the spectroradiometric measurements of cholesteatoma tissue using a linear combination of the fluorescence emitted by keratin and FAD powder measured under the same illumination and measurement conditions.

The presence of keratin is one of the signature characteristics of cholesteatoma. Cholesteatoma is composed of a matrix, perimatrix and cystic content.^19^ The cystic content, is composed of anucleated and fully differentiated keratin scales and necrotic material.^40^ This keratin content is highly fluorescent as seen in Figure 6. The matrix is made up of highly metabolically active stratified squamous epithelium hyper-proliferating and producing keratin lamellas that reach the cystic part. The perimatrix is sub-epithelial connective tissue that contains collagen fibers, elastic fibers, fibroblasts and inflammatory cells. The perimatrix has less overall fluorescence than that of the matrix and cystic content, as shown in Figure 11. Therefore, the keratin from the cystic content and the metabolic activity of the matrix can explain the high keratin and FAD signal we measured in the autofluorescence spectra.

The ability to excite more than one fluorophore is clinically important since identification does not rely solely on keratin fluorescence but also on metabolic activity due to FAD. FAD is a coenzyme involved in the cell metabolism, similar to NADH. NADH and FAD fluorescence have been previously used to provide information regarding the tumor micro-environment and can be used to noninvasively monitor changes in metabolism.^41, 42^ Additionally, through our microscope slide images, it is likely that the largest components of keratin are located in pearls, as shown in Figure 11a-c. In this case, even though the keratin autofluorescence signal is larger than FAD (based on Figure 6, keratin may not be found at every location of the cholesteatoma tissue. Looking at Figure 11c the areas of 405 nm fluorescence shown in green and 450 nm fluorescence shown in red are not the same locations. So, it is still essential to utilize the 450 nm wavelength for FAD because not only does it provide information on the environment and metabolic activity, the FAD signal also allows for a more complete visualization of the tissue. As shown in Figure 11c, the two fluorophores of FAD and keratin are microscopically located in different areas. The keratin component excited by the 405 nm illumination is represented in green and primarily exists in the folds of the keratin pearl. In contrast, the FAD component excited by 450 nm illumination and shown in red, fluoresces more strongly in between the folds. By utilizing both wavelengths, the device will be able to pick up the clearest and most complete visual of the cholesteatoma tissue.

Additionally, in many cholesteatoma operation cases, there is the presence of bacteria such as *Pseudomonas aeruginosa* and *Staphylococcus aureus* surrounding the cholesteatoma tissue.^43, 44^ These types of bacteria, along with many others, contain porphyrins which also have a very prominent auto-fluorescent signature.^45^ Considering this, the cholesteatoma spectra measured in Figure 4 show a slight fluorescent signal in the characteristic location of porphyrin autofluorescence (650 nm to 700 nm).^46^ While the signal is very faint, there could be a small amount of bacteria on these samples. Furthermore, when the device is used in the operating room, it is important to consider the possibility and likelihood of bacteria being present during the operation. Therefore, in those cases, our device could take advantage of the porphyrin emission using our 405 and 450 nm excitation illumination to visualize the presence of bacteria.

These results motivated the design of an imaging system based on 405 and 450 nm illumination and a camera equipped with a 495 nm longpass filter that can be adapted to the clinic or operating room. Therefore, our imaging system would only add the addition of fluorescent imaging on top of the current gold standard white light imaging, so surgeons can switch between our newer imaging format and what they are already familiar with. We captured camera images of cholesteatoma and mucosa tissue using the same illumination and physical conditions that were used for the spectroradiometric measurements (Figure 4). Both measurements show that cholesteatoma tissue samples fluoresce when illuminated with 405 and 450 nm light, and mucosa tissue samples do not.

We compared the measured camera RGB pixel values for cholesteatoma tissue with predicted camera RGB pixel values that were obtained by multiplying the spectrophotometric measurements of cholesteatoma tissue with the camera quantum efficiency (QE). Our understanding of the impact of the camera components on measuring cholesteatoma tissue fluorescence is validated by the agreement between the predicted and actual camera values, demonstrating the camera’s ability to produce accurate results.

The light that is emitted from tissue (i.e. tissue autofluorescence) is weak relative to the light that is reflected from tissue. Although we used a longpass filter to block reflected light, tissue autofluorescence is still a very weak signal. In order to obtain camera images of tissue autofluorescence with high SNR, we used a relatively long camera exposure duration (500 ms). Since the tissue did not move during the exposure duration, the camera images we captured were not blurred. However, if tissue moves (such as in surgery), camera images will be blurred. There are several ways to minimize or even eliminate motion blur and still capture images with high SNR. For example, once can reduce the exposure duration by either increasing the intensity of the light (fluorescence is linear with the intensity of the excitation light) or by increasing the camera sensitivity (i.e. quantum efficiency) to the emitted light. Another design option is to fix the camera position in such a way that movement of the camera is matched (yoked to) movement of the tissue.

In future research, we plan to explore these design options and evaluate the imaging system *in vivo* in an animal model. We are able to induce cholesteatoma^47, 48^ in gerbils by ligating their Eustachian tubes. Since blood is always prevalent in any surgical setting, there is a likelihood that the autofluorescence signal of the cholesteatoma will be diminished by light absorption at 545 and 580 nm.^49^ For example, we observed dips in the spectroradiometric measurements of cholesteatoma fluorescence at 545 and 580 nm that we observed (see Figure 4b). This effect was very small in this study, but our *in vivo* testing will give us a better understanding of how the presence of blood can affect the measurement of cholesteatoma tissue fluorescence.

## 5. CONCLUSION

We were able to leverage the autofluorescence of cholesteatoma to design an imaging system composed of a light source, filters, and a camera that can provide image-guided assistance during the surgical removal of cholesteatoma. We were able to accurately predict the device images using the camera QE, validating that the spectral measurements and the device images are consistent with one another.

## 7. DATA AVAILABILITY

All data in support of the findings of this paper are available within the article.

## ACKNOWLEDGMENTS

We thank our collaborators from Stanford Medicine Children’s Health, Department of Otolaryngology - Head & Neck Surgery: Nikolas Blevins, MD; Kay Chang, MD; and Alan Chang, MD; for contributing to cholesteatoma and mucosa sample collection from the operating room.

